# Analysis of Biological Noise in an Organelle Size Control System

**DOI:** 10.1101/2020.08.31.276428

**Authors:** David Bauer, Hiroaki Ishikawa, Kimberly A. Wemmer, Jane Kondev, Wallace F. Marshall

**Affiliations:** Dept. of Biochemistry & Biophysics University of California, San Francisco 600 16th St., San Francisco, CA; Dept. of Physics Brandeis University Abelson-Bass-Yalem Building, 97-301, Waltham, MA; Center for Cellular Construction

## Abstract

Analysis of fluctuation in organelle size provides a new way to probe the mechanisms of organelle size control systems. By analyzing cell-to-cell variation and within-cell fluctuations of flagellar length in *Chlamydomonas*, we show that the flagellar length control system exhibits both types of variation. Cell to cell variation is dominated by cell size, while within-cell variation results from dynamic fluctuations that are subject to a constraint, providing evidence for a homeostatic size control system. We analyzed a series of candidate genes affecting flagella and found that flagellar length variation is increased in mutations which increase the average flagellar length, an effect that we show is consistent with a theoretical model for flagellar length regulation based on length-dependent intraflagellar transport balanced by length-independent disassembly. Comparing the magnitude and time-scale of length fluctuations with simple models suggests that tubulin assembly is not directly coupled with IFT-mediated arrival and that observed fluctuations involve tubulin assembly and disassembly events involving large numbers of tubulin dimers. Cells with greater differences in their flagellar lengths show impaired swimming but improved gliding motility, raising the possibility that cells have evolved mechanisms to tune intrinsic noise in length. Taken together our results show that biological noise exists at the level of subcellular structures, with a corresponding effect on cell function, and can provide new insights into the mechanisms of organelle size control.

## Introduction

The mechanisms by which cells control the size of their organelles remain incompletely understood. Indeed, defining what it means for organelle size to be “controlled” is not entirely clear. In some cases, limited availability of molecular precursors can limit organelle size (Goehring 2012), but such a limiting precursor condition is unlike what we normally think of as “control”, where active systems monitor a quantity and adjust it back to a pre-defined set point. One way that a control system can manifest itself is by constraining variation. In the case of organelles, this viewpoint suggests that the function of a size control system could be explored by examining the variation in organelle size within or among cells. Analysis of biological fluctuations (noise) have been extensively employed in neurobiology to probe mechanisms regulating single-channel opening and closing (Neher and Stevens, 1977) as well as in the study of gene expression (Elowitz et al, 2002; Raser et al, 2004; Kaern et al., 2005; Perkins and Swain. 2009). An important contribution of gene expression noise studies was to draw attention to the difference between “intrinsic” noise, caused by fluctuations within the machinery of a promoter at each individual gene, for example due to stochastic binding of transcription factors, and “extrinsic” noise, caused by variation in the rest of the cell that affect all genes, for example variation in the number of RNA polymerase molecules, Cell to cell variation in organelle size and number have been observed even in genetically identical cells (Marshall 2007; Mukherji 2014; Chang 2017; Chang 2019). For example, analysis of variation in the copy number of various organelles, as well as the joint variation in organelle number and size, have been used to gain insight into the mechanisms of organelle dynamics (Mukherji 2014; Amiri 2019).

Here we describe both cell-to-cell “extrinsic” variation and within-cell “intrinsic” fluctuations in the size of flagella, a model organelle chosen for its simple geometry, which allows size to be defined by a single number, the length. Cilia and flagella are interchangeable terms for the microtubule-based organelles that project from the surface of most eukaryotic cells (Pazour and Rosenbaum, 2002) and perform important motile and sensory functions (Scholey and Anderson, 2006). Because these organelles are simple linear structures, it is easier to analyze the length regulation of cilia and flagella than it is to analyze size control in more complicated organelles. The flagellum has proven to be an excellent test-bed for exploring the systems biology of cellular structure (Randall, 1969; Wemmer and Marshall, 2007). The lengths of cilia and flagella are known to vary from cell to cell in a variety of organisms (Randall, 1969; Wheatley and Bowser, 2000; Adams et al., 1985) suggesting these organelles can be used to analyze biological noise at the level of cellular structure. The length distribution of flagella is not Gaussian, and different mutants with similar average lengths can be distinguished on the basis of differences in the shape of the length distribution (Kannegaard 2014), supporting the idea that length variation is an important aspect of the length phenotype. However, previous experimental studies of flagellar length variation have not distinguished between sources of variation that take place within the flagella (intrinsic noise) versus source of variation that take place in the cell body (extrinsic noise). Theoretical models for flagellar length regulation make distinct predictions concerning the nature of flagellar length fluctuation and variation (Bressloff 2006; Hendel 2018; Fai 2019; Banerjee 2020; Patra 2020), hence a measurement of length fluctuation has the potential to test existing models and inform future modeling efforts.

Here, we use measurements of flagellar length to measure two components of flagellar length variation: a correlated component that reflects cell-to-cell variation and an uncorrelated component that reflects within-cell variation in length between the two flagella. We demonstrate that these types of variation stem from a combination of intrinsic and extrinsic biological noise. The existence of an active length-control system is indicated by a constraint on fluctuations in individual flagella. We leveraged *Chlamydomonas* genetics to explore molecular pathways that might contribute to noise, showing that mutants increasing length lead to an increase in the uncorrelated component of variation, reflected by an increase in the magnitude and time scale of length fluctuations. This result is then shown to be consistent with predictions of a linear noise model derived from a previously published flagellar length control model. Finally, we show that differences in flagellar length have an effect on biological fitness in terms of cell motility, and suggest that the biological noise seen in wild-type vegetative cells may represent a balance between the requirements of two different types of motility.

## Results

### Measuring biological noise in the flagellar length control system

In order to analyze noise in flagellar length, we measured lengths in the unicellular green alga *Chlamydomonas reinhardtii* (Merchant et al., 2007), a well-established model organism for the study of cilia and flagella. Each *Chlamydomonas* cell has two flagella approximately 10 μm long (**Figure 1A** inset). This organism has long been used in studies of flagellar length control because its flagella are easy to measure and because mutants exist in which flagellar length is altered (Wemmer and Marshall, 2007).

**Figure 1.**
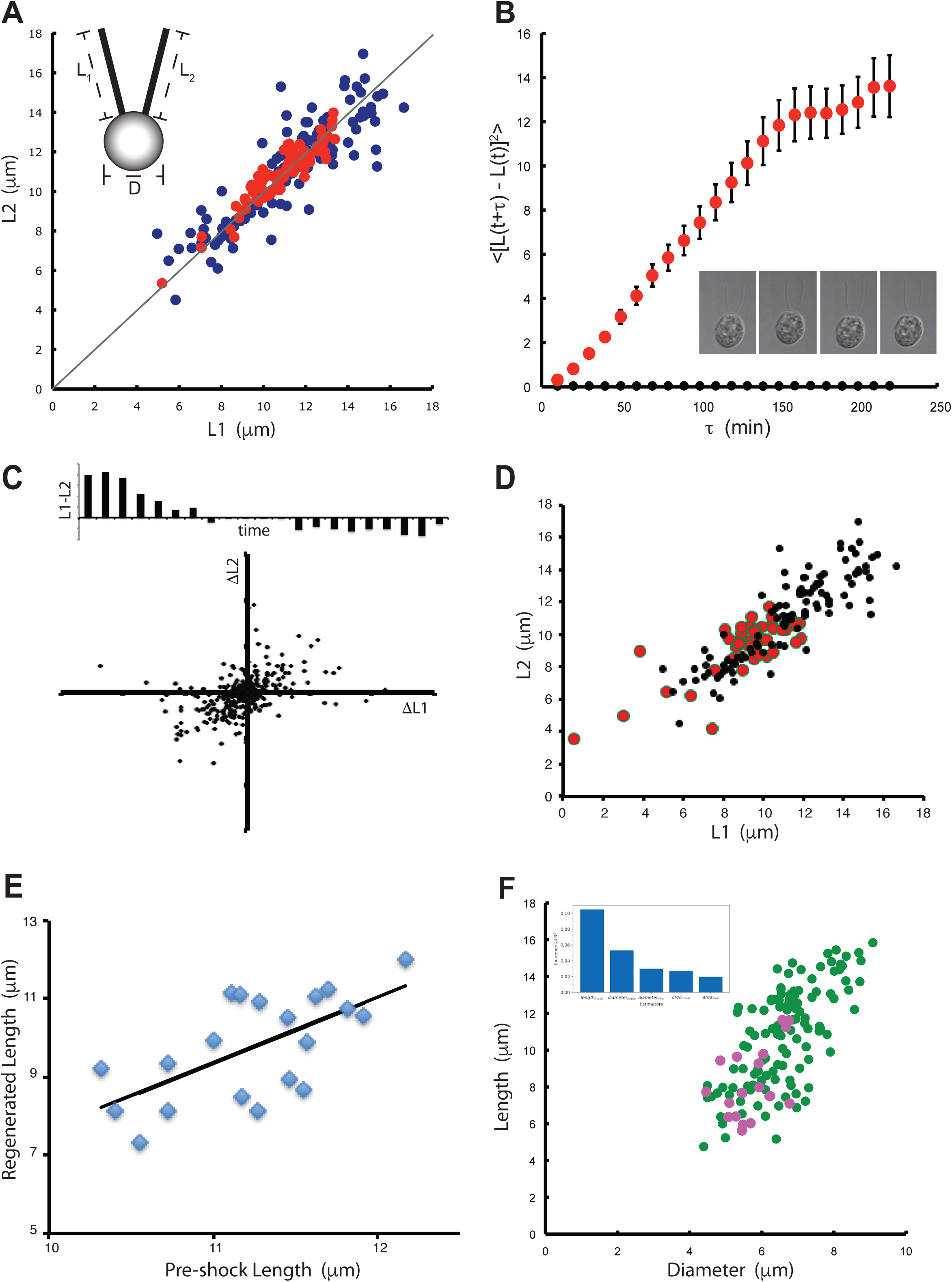
Measuring intrinsic and extrinsic noise in flagellar length control system. **(A)** Intrinsic and extrinsic noise can be visualized in measurements of flagellar length in fixed *Chlamydomonas* cells with two equivalent flagella. Graph plots length of one flagellum versus length of the other flagellum. Extrinsic noise is reflected by scatter along the diagonal L_1_=L_2_ (gray line). Intrinsic noise is reflected by scatter perpendicular to this axis. (Blue) wild-type asynchronous culture, (Red) wild-type gametes. (Inset) Cartoon of a *Chlamydomonas* cell showing the three measurements reported in this paper. (**B**) Fluctuations in length observed in living cells. (Red) Mean squared change in length plotted versus time, showing constrained diffusion-like behavior. Error bars signify standard error of the mean. (Black) Mean squared change in length in glutaraldehyde-fixed cells as an indicator of measurement error. (Inset) Four successive time points, taken ten minutes apart, of a 3D time series of a single living cell embedded in agarose and imaged with DIC microscopy, similar to the images used to generate the data in this plot. (**C**) Joint fluctuations in two flagella. Lower plot: scatter plot showing fluctuations in the lengths of both flagella during a ten-minute time-step. Upper plot: length difference between two flagella in one cell, plotted for successive time-points, showing signal reversal, indicating that neither flagellum is consistently longer than the other. (**D**). Flagellar length variation in *ptx1* mutant that eliminates functional asymmetry between the flagella (red). Wild-type data from Panel A is reproduced for reference as black circle. (**E**) Flagellar length before and after regeneration induced by pH shock in a microfluidic device. Line indicates best fit to data. **(F)** Contribution to correlated length variation from cell size variation measured by correlation between average length of the two flagella with the cell diameter in asynchronous cultures. (Green) Wild-type cells. (Pink) *mat3* mutant cells that have smaller average size than wild-type (Umen and Goodenough, 2001) were used to extend the available size range for analysis. Correlation coefficients were 0.61 for wild-type cells alone and 0.70 for the combination of wild-type cells plus *mat3* cells. Inset gives result of dominance analysis of flagellar length after regeneration (based on data in Supplemental Figure S1E in terms of length prior to regeneration and different measures of cell size. Plot shows the incremental r^2^ contribution of each predictor in the model.

Our quantitative analysis of noise in flagellar length is based on approaches previously developed for gene expression analysis. Two distinct types of noise have been described for studies of gene expression: extrinsic noise, which describes the variation in overall expression levels from cell to cell, and intrinsic noise which describes variations in transcription rates at an individual promoter (Elowitz et al, 2002; Raser et al., 2004). In gene expression studies, the distinction between the two types of noise is measured using a dual-reporter method (Elowitz et al., 2002), in which two different fluorescent protein reporter constructs are driven by identical promotors, so that variation in expression of both constructs can be measured on a cell-by-cell basis. The ability to distinguish between fluctuations within a single gene from variation that is shared by the whole cell has been critical to understanding the mechanisms and functional implications of transcriptional noise. In applying these concepts to flagellar length control, we consider the “system” to be the flagellum, of where there are two copies as in a dual reporter gene construct, and the “environment” to be the cell body along with any factors outside of the cell. We choose this way of decomposing the cell because we are specifically interested in control mechanisms that may exist within flagella to constrain length variation.

The mathematical decomposition of noise sources into intrinsic and extrinsic, achieved by decomposing total variation from dual-reporter experiments into correlated and uncorrelated components, has been called into question on the grounds that it relies on numerous assumptions that may not hold for a system of interest (Hilfinger and Paulsson, 2011). For example, it is assumed that noise sources are additive, which may not be the case. Of particular relevance to flagellar length control is the fact that the intrinsic/extrinsic decomposition breaks down in the situation where the two systems are competing for a common resource (Stamatakis 2011), which is the case for flagella, which compete for a common cytoplasmic pool of precursor proteins. Based on such considerations, it is potentially misleading to equate the correlated and uncorrelated components of total variance with extrinsic and intrinsic sources, respectively. Nevertheless, the decomposition is still useful because it highlights two types of variation that are expected to have distinct biological impacts – namely the variation in length of the two flagella within one cell versus the variation in the lengths of both flagella from one cell to another. These types of variation may reflect different underlying processes, and may have different effects on cellular function. We will refer to these two components of total variance as the uncorrelated and correlated noise components, respectively, where the correlation in question is that between the two flagella in a single cell. Later, once we have described the measurements of these components of variation, we will address their possible relation, if any, to actual intrinsic or extrinsic sources.

We are able to measure correlated and uncorrelated noise in the case of flagellar length, by exploiting the fact that *Chlamydomonas reinhardtii* cells naturally have two identical flagella. We measured the lengths of both flagella in a population of cells (see **Figure 1A**) and from these data we computed the correlated and uncorrelated noise components, using the same mathematical formalisms used for gene expression studies (See Materials and Methods), essentially treating the two flagella as two reporter genes in a dual-reporter experiment.

As shown in **Figure 1A**, and summarized in **Table 1**, flagellar lengths in wild-type cells demonstrated measurable levels of both correlated (variation in L1 – L2) and uncorrelated noise (variation in L1 + L2 – Lavg). Uncorrelated noise, reflected by the deviation of points away from the diagonal, describes the variations in length between the two flagella within one cell and may represent, at least in part, fluctuations within the length-control machinery intrinsic to each flagellum. Correlated noise, reflected by the variation of points along the diagonal, describes the cell to cell variation in length and may represent, at least in part, fluctuations in cell-wide features such as cell size (see below). We found that the correlated noise is greater in magnitude than the uncorrelated noise. The uncorrelated noise is, however, significantly greater than the estimated measurement error (see Materials and Methods), thus we can conclude that flagella are subject to measurable biological noise.

**Table 1.**
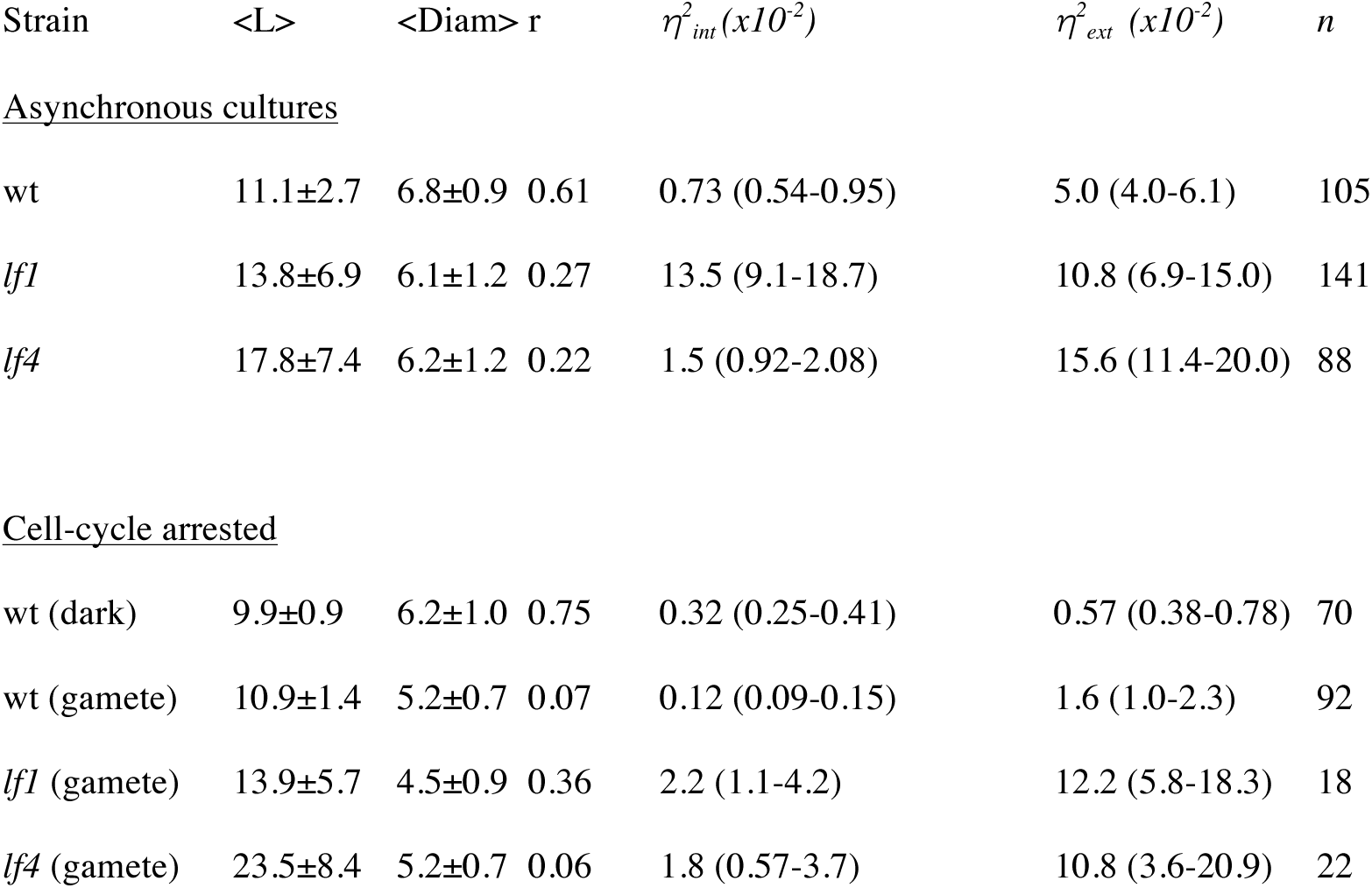
Summary of noise calculations. <L> denotes average flagellar length in μm, with standard deviation listed. <Diam> is the average cell diameter in μm. r is the correlation coefficient between flagellar length and cell diameter. *η^2^* and *η^2^* denote the uncorrelated and correlated components of variation, respectively. Noise measures η^2^ listed are the actual measurements multiplied by 100. n is the number of cells measured. 95% confidence intervals for noise measures are given in parentheses.

### Constrained fluctuation in flagellar length reveals a length control system

**Figure 1A** depicts variation in flagellar length in a population of fixed cells, and raises the question of what is the source of this variation. Does this variation reflect stable differences between flagella, or randomly fluctuating processes? Perhaps the simplest basis for flagellar length variation might be that when flagella are first formed, they assemble in a range of lengths and then maintain them in a static manner. This sort of baked-in size variation is often seen in quenched physical systems, for example grain size variation in metallurgy when a metal is rapidly quenched and can be seen in biological examples such as bacteriophage tail lengths, which assemble in a range of sizes but, once assembled, do not undergo any further length changes. To ask whether flagella length variation is fixed or dynamic, we embedded cells in agarose and acquired time-lapse 3D movies using DIC microscopy, allowing us to measure length fluctuations as a function of time. A plot of the mean-squared change in flagellar length as a function of time-interval (**Figure 1B**; plot shows calculated mean-squared changes for both individual flagella averaged in multiple cells as described in Materials and Methods) shows that flagellar lengths appear to undergo random-walk dynamics at short time scales, with a mean squared length change linearly proportional to the time interval. At longer time-scales, the mean squared length difference reaches a plateau, suggesting that the magnitude of the length variation is constrained. This constraint on length variation presumably reflects the existence of a flagellar length control system. This live-cell analysis demonstrates that flagellar length variation is not simply locked in from the time of initial assembly, but rather reflects an ongoing series of fluctuations. When fluctuations in the two flagella are compared in single cells (**Figure 1C**), we find that the fluctuations show a small but statistically significant correlation (r=0.32). One possible explanation for this result could be that the two flagella communicate with each other, such that noise processes occurring in one flagellum influence fluctuations in the other. One possible mechanism for communication between the two flagella is competition for a shared limiting precursor pool (Kuchka and Jarvik 1982) such that if one flagellum becomes longer, the other would be forced to become shorter. Such competition would be expected to produce an anticorrelation, unlike the positive correlation we observe. The positive correlation thus appears to suggest that competition for a shared pool is not a major determinant of length variation in our experiments. An alternative explanation for the correlated fluctuations is that noise processes in the cell body may drive length fluctuations in both flagella.

### Evidence for extrinsic and intrinsic noise sources in flagellar length control system

Can the uncorrelated and correlated parts of the variation in flagellar lengths be ascribed to intrinsic noise or to extrinsic noise, respectively? These two types of noise are defined relative to a system of interest, in this case, the flagellum. Intrinsic noise, in this context, refers to variation in flagellar length resulting from variations in biological processes taking place entirely within the flagellum, such as intraflagellar transport or tubulin polymerization at the flagellar tip. Extrinsic noise, in this context, refers to variations taking place outside of the flagellum, specifically, process in the rest of the cell, which serves as the biochemical environment within which the flagellum assembles and is maintained. Examples of extrinsic sources of variation would include flagellar protein synthesis and intracellular transport of cargo and motors to the base of the flagella. Do both types of noise exist for flagella? Given the number of assumptions required to equate correlated and uncorrelated variation with extrinsic and intrinsic noise, we cannot infer the presence of both types of noise just on the basis of our measurements of variance. Instead, we seek to test for intrinsic and extrinsic noise via direct experiments.

Intrinsic noise sources within the flagellum would result in uncorrelated variation in their length, however we must consider the alternative: that the uncorrelated variation is not caused by flagella intrinsic factors. One might imagine that when two flagella are first built, there is some difference in length that is then permanently locked in. We have ruled out locked-in length variation by showing that flagellar lengths fluctuate (**Figure 1B**). However, suppose some varying process in the cell body such as precursor production had an influence on flagellar length, and the two flagella had some locked-in differences in their ability to respond to this input (for example, if they differed in the quantity of IFT machinery required to transport precursor proteins in each flagellum). In such a case, the lengths of both flagella would fluctuate in response to the extrinsic fluctuation in the cell body, but they would respond to that extrinsic fluctuation through different multiplicative factors, such that the difference in their length would itself fluctuate, giving rise to an uncorrelated component of the total variation. In such a scenario, although both flagella would change length, the longer of the two flagella would always be longer and the shorter one would always be shorter, since the difference in length reflects an underlying static difference in their ability to utilize precursor. We therefore analyzed the signed difference in length between the two flagella and asked if it undergoes sign changes. An example is shown in **Figure 1C**, in which it is clear that the flagellum that was initially longer, ends up being shorter at the end of the time series. Out of sixteen live cells analyzed, eleven show at least one sign change in the difference between the flagellar lengths during the course of observation. This result indicates that the uncorrelated variation is unlikely to be due to a multiplicative effect of extrinsic fluctuations.

We also tested one concrete mechanism for locked-in variation in the two flagellar length control systems, based on the fact that the two flagella of a *Chlamydomonas* cell have subtle functional differences that derive from the age of the basal bodies at the base each flagellum (Kamiya and Witman 1984). These functional differences, which are required for cells to steer towards a light source during phototaxis, are eliminated in the phototaxis deficient *ptx1* mutant (Horst 1993). We therefore measured the uncorrelated and correlated components of variation in *ptx1* mutants versus wild-type. If asymmetry in flagella caused a consistent variation in length, we would expect the uncorrelated variation to be reduced in this mutant. As shown in **Figure 1D**, although the average length in *ptx1* mutants is reduced compared to wild type, the uncorrelated variation is larger than in wild type *(η^2^*_*int*_ = 0.013 [95% confidence interval 0.007 – 0.02] for *ptx1* versus *η^2^*_*int*_ = 0.007 [95% confidence interval 0.005 – 0.01] for wt). We conclude that the PTX1-dependent biochemical asymmetry in the two flagella is not a substantial source of uncorrelated variation. Based on these experiments, we conclude that at least some of the uncorrelated variation results from intrinsic fluctuations within the flagella.

We next asked whether some of the correlated variation could be driven by extrinsic noise, i.e., by variation in the cell body. To test for extrinsic noise directly, we took advantage of the fact that *Chlamydomonas* cells can rapidly regrow their flagella after detachment, which is easily triggered by transient pH shock (Quarmby 1994). We used two microfluidic device, one commercial and one custom-built, to trap individual *Chlamydomonas* cells and hold them in position while media is exchanged (see **Supplementary Figure S1A and C**). Using these systems, we imaged flagella in single cells before, during, and after regeneration, allowing us to ask whether cells with long versus short flagella would regenerate correspondingly long or short flagella after pH shock. Because flagella are completely removed during this experiment and regrown, any correlation in length before and after regeneration would have to reflect a property of the cell body itself. As shown in **Figure 1E and Supplementary Figure S1B and E**, there is a significant correlation between average flagella length before deflagellation and after regeneration (r=0.65; P=0.0011 for data in **Figure 1E**). We note, in these experiments, that flagella are shorter after regeneration compared to their pre-pH shock length. This observation is consistent with previous studies of flagellar regrowth (Rosenbaum 1969), which show that while flagella rapidly grow to within a micron of their final length, they then continue to grow at a much slower rate for many hours. The high correlation observed between length before pH shock and length after regeneration shows that flagella are subject to extrinsic variation due to differences in the cell body of different cells. Because the flagella are removed by the pH shock, the determinant of length must reside with the cell body. What property of a cell drives this variation?

Extrinsic noise in gene expression is dominated by variation in cell size (Raser et al., 2004), presumably because larger cells have more ribosomes thus making more protein per mRNA molecule. Flagellar length depends on the availability of flagellar precursor proteins (Rosenbaum et al., 1969; Coyne and Rosenbaum 1979), suggesting that larger cells might have longer flagella due to increased numbers of ribosomes. Indeed, it has been noted that larger cells sometimes have longer flagella in various mutants (Adams et al., 1985). We found that the correlated component of flagellar length variation, as quantified by the average of the lengths of the two flagella in a given cell, was strongly correlated with cell size (**Figure 1F**). Dominance analysis of flagellar length before and after regeneration in our pH shock experiments (inset **Figure 1F**) indicates that cell size is one predictor of flagellar length after regeneration, but that length prior to regeneration is a stronger predictor, indicating that other features of cell state must affect length besides just cell size. The other obvious candidate is cell cycle state. We compared flagellar length variation in asynchronous cultures (which were used to obtain the data discussed above) to flagellar length variation in cells arrested in G1 by growth in the dark in minimal media, as wells as to gamete cultures which are arrested in a G0-like state in preparation for mating. Correlated variation was significantly reduced by both types of cell cycle arrest (**Table 1**), suggesting that cell-to-cell variations in cell cycle state normally contribute to correlated variation in the asynchronous cultures. As shown in **Table 1**, the correlation between cell size and flagellar length is stronger in G1 arrested cells (r=0.75) than in asynchronous cultures (r=0.61) suggesting that when variation due to cell cycle progression is eliminated, the remaining correlated variation is more closely determined by cell size variation. Because cell size and cell cycle state are features of the cell body, the fact that these two features influence the correlated variation in flagellar length suggests that this correlated variation is indeed due to extrinsic noise, i.e., to random variation in processes taking place in the cell body.

### Identifying genes that regulate noise in flagellar length control system

Fluctuation analysis provides a tool for probing control systems by exploring how various components of a system alter the observed fluctuations. We used this approach to test a series of candidate genes for possible roles in reducing uncorrelated variation in flagellar lengths. Mutations that increase uncorrelated variation could potentially reveal genes involved in constraining flagellar length variation or coordinating lengths of the two flagella within a cell. We therefore tested a panel of mutants affecting a range of cellular processes for possible roles in noise suppression (**Figure 2A**). First we asked whether transcription or translation were needed to actively regulate uncorrelated noise. In cells treated with cycloheximide as previously described (Rosenbaum et al., 1969), uncorrelated noise was not significantly altered compared to untreated cells, indicating that variation in gene expression is unlikely to play a role in regulating flagellar length on a dynamic basis.

**Figure 2.**
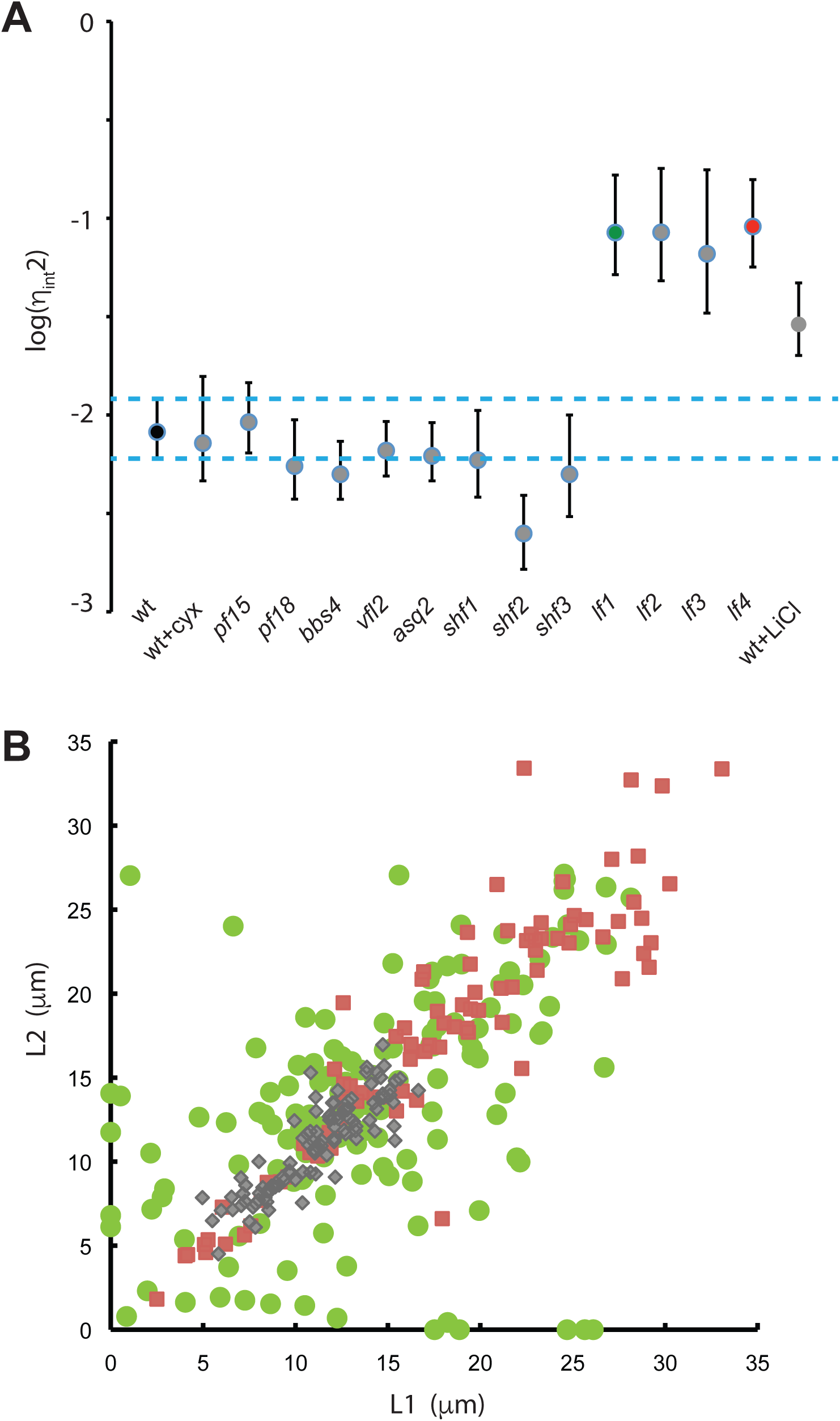
Identification of genes that participate in flagellar length noise suppression. (**A**) Analysis of candidate genes for effect on noise. Points represent median calculated value of uncorrelated variation. Error bars are 95 percent confidence intervals for each strain. Dotted blue lines show the 95 percent confidence interval for uncorrelated variation in wild-type cells. Green and Red data markers indicate *lf1* and *lf4* mutants. All measurements were made using 2D image data to allow a large number of different strains to be measured more rapidly. (**B**) Increased correlated and uncorrelated variation in long flagella mutants. (Green circles) *lf1*, (Red squares) *lf4*, (Gray diamonds) wild-type. All data shown is from 3D measurements of flagellar length in cells from asynchronous cultures.

We next considered possible mechanisms for communication between flagella that might play a role in constraining uncorrelated variation. One way that flagella communicate with each other is via hydrodynamic coupling between the two beating flagella (Brumley 2014). We examined paralyzed mutants *pf15* and *pf18* in which flagella are non-motile, and found they had no increase in uncorrelated variation.

A second way that flagella communicate is via protein fibers connecting their basal bodies, which are known to play a role in synchronization of flagellar beating (Wan 2016). We therefore analyzed *asq2* mutant cells, which we have previously shown to be defective in the physical connections between mother and daughter basal bodies (Feldman 2007; Feldman 2009), and *vfl2* mutants which lacks fibers that normally connect the basal bodies to the nucleus (Wright et al., 1985). Neither mutation had an effect on uncorrelated variation.

We next examined mutants with altered flagellar length. Mutations in the SHF1, SHF2, and SHF3 genes have been reported to cause cells to form short flagella, roughly half wild-type length. Noise measurements in these mutants did not show any increase in uncorrelated variation. In contrast, we saw a dramatic increase in uncorrelated variation in *lf1*, *lf2*, *lf3*, and *lf4* mutants, all of which have abnormally long flagella (McVittie, 1972; Barsel et al., 1988; Berman et al., 2003; Nguyen et al., 2005; Tam et al., 2007; Wemmer 2019). This suggested that increased length might somehow lead to increased uncorrelated variation. To confirm this hypothesis, we measured noise in wild-type cells treated with lithium, which increases flagellar length (Nakamura et al., 1987) due to an increase in IFT activity (Ludington 2013). As with the *lf* mutants, we found that lithium treated cells also showed increased uncorrelated length variation.

To further examine the role that LF genes play in noise regulation, we examined individual mutants in more detail. **Figure 2B** plots the distribution of flagellar length pairs in *lf1* and *lf4* mutants. Both mutants cause the average flagellar length to be approximately double wild-type length. It was previously reported that the *lf1* mutation leads to increased variance in length (McVittie, 1972) but in that analysis no attempt had been made to distinguish intrinsic and extrinsic variation. As indicated in **Figure 2B**, we found that both mutations lead to increased noise, but in different ways: *lf1* mainly increases uncorrelated variation while *lf4* increases both types of variation (**Table 1**). One obvious difference is that *lf1* mutants are more likely to be uniflagellar, which is one of the reasons that the uncorrelated variation is higher in *lf1* than *lf4*, but even ignoring such cells, it is clear from **Figure 2B** that *lf1* show greater differences between the lengths of their two flagella than *lf4* mutants. This suggests the two genes act in fundamentally different ways within the length control system. For both mutants, the quantitative effect of the *lf* mutations on noise is much greater than their effect on average length (Table 1), suggesting that the LF gene products may have evolved primarily as a means to restrain length variation, as opposed to setting any particular average length.

Given that correlated variation appears to result from extrinsic noise (**Figure 1E,F**), one possible explanation for increased uncorrelated variation in *lf4* mutants could be that *lf4* mutants have a wider variation in cell size. Indeed, the standard deviation in cell diameter in *lf4* is larger than wild-type cells by a ratio of 1.3. However, the increase in cell size is far less than the increase in standard deviation in flagellar lengths, so increased cell size variability is unlikely to be the sole explanation for the increased extrinsic noise in *lf4*. The other possibility is that the *lf4* mutation may cause length to depend more strongly on cell size than in wild-type. This possibility is supported by the fact that the slope of the best-fit line relating flagellar length to cell diameter increases from 0.96 in wild type to 1.74 in *lf4* cells. We speculate that this difference may lie in the fact that *lf4* mutants show an increased rate of intraflagellar transport (Ludington 2013; Wang 2019), so that a given quantity of precursor protein such as tubulin can be more efficiently used for flagellar growth.

We conclude from this limited candidate screen that LF1 and LF4 genes act to reduce biological noise in flagellar length. In fact, both genes seem to have a larger effect on noise than they do on the average flagellar length, suggesting that noise suppression may be their primary evolutionary purpose.

### Noise analysis of balance-point model for flagellar length control

The noise measurements reported here provide a new way to test hypothetical flagellar length control system models, since any model for such a system must be able to account for not only the average steady-state behavior of the system, but also the fluctuations that occur within the system. For example, any model that claims to explain flagellar length control must be able to account for the observation presented above that mutations leading to increased average length also lead to increased uncorrelated variation.

We previously described a model, outlined in **Figures 3A and B**, for flagellar length control (Marshall and Rosenbaum, 2001; Marshall et al., 2005), based on the intrinsic length-dependence of intraflagellar transport (IFT). IFT is mediated by the kinesin-driven movement of protein complexes called IFT particles which bind to and transport tubulin and other flagellar building blocks out to the flagellar tip, where they assemble into the growing structure (Qin et al., 2004; Hao et al., 2011; Bhogaraju et al., 2013; Ishikawa, 2017). Our IFT-based length control model contains three parameters, A, D, and P, which represent the efficiency of IFT, the rate of disassembly at the tip, and the level of flagellar structural protein synthesized by the cell. As previously derived (Marshall and Rosenbaum, 2001) and explained further in Materials and Methods, the steady-state average flagellar length for this model is:

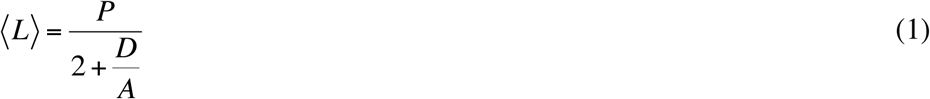

**Figure 3.**
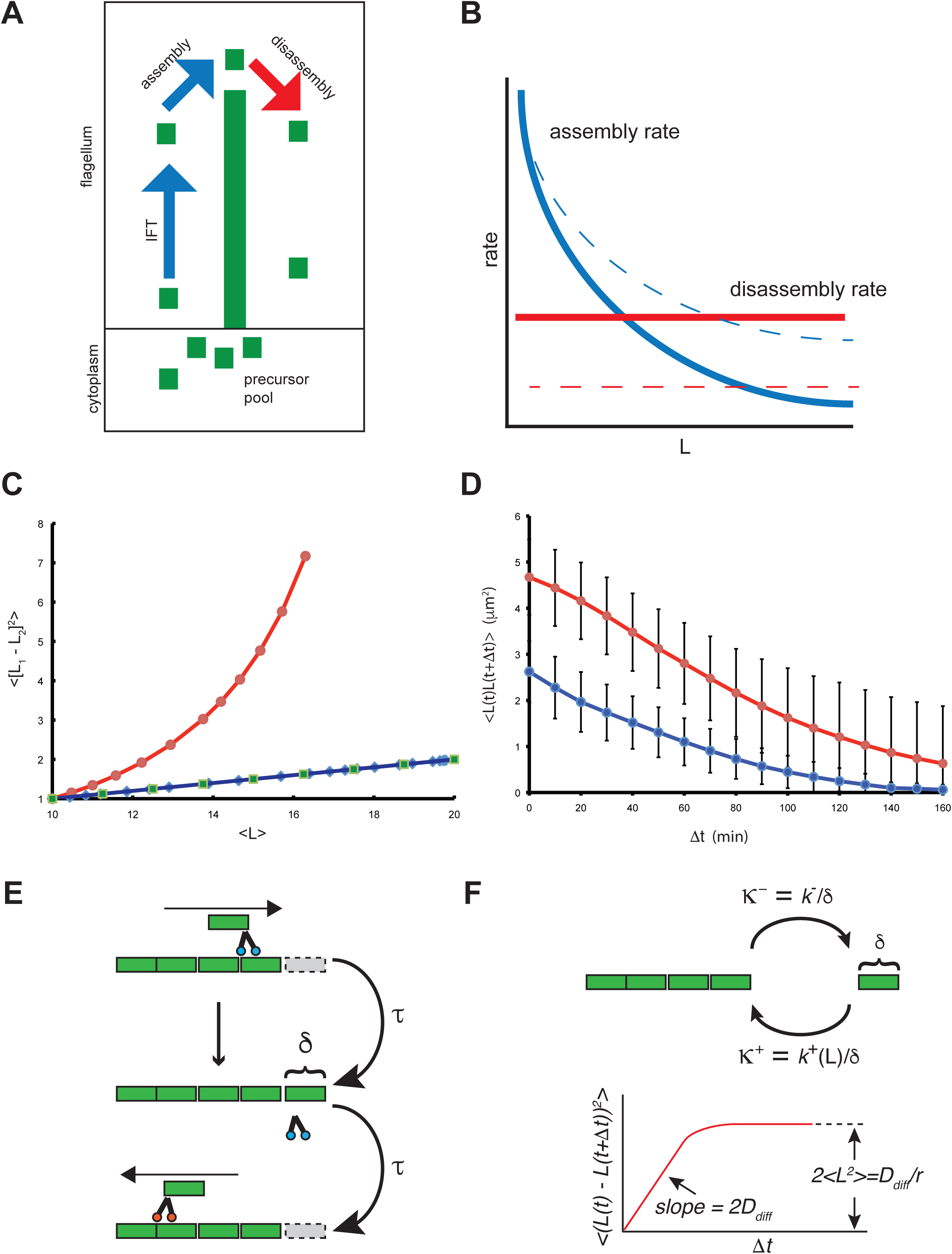
Linear noise analysis of a length control model. **(A)** Schematic of balance-point mode for flagellar length control. Flagellar microtubules undergo constant disassembly (red arrow) at a length-independent rate D. This disassembly is balanced by assembly, which occurs at a rate limited by the rate of IFT (blue arrow). Assuming the number of IFT particles is length-independent, the rate of cargo delivery at the tip is proportional to 1/L. Precursor protein from the cytoplasm binds with first order binding. The available free precursor pool is the total pool P minus the quantity of precursor already incorporated into the two flagella. Hence the net assembly rate is given by A(P-2L)/L. (**B**) Steady-state length is determined by the balance point between length-independent disassembly (Red line) and length-dependent assembly (Blue line). Mutations can increase flagellar length either by increasing assembly (Blue dotted line) or decreasing disassembly (Red dotted line). **(C)** Results of small signal noise analysis assuming an intrinsic noise source acting within one flagellum. Plot shows the mean squared difference in the lengths of the two flagella plotted versus their average length, while parameters A (describing the efficacy of IFT), D (describing the rate of steady state axoneme disassembly), or P (describing the total pool of flagellar precursor protein) are changed so as to increase the steady-state length from 10 to 12 microns. (Red circles) results of decreasing D, (Blue diamonds) results of increasing A, (Green squares) results of increasing P. Since the strength of the noise source is not known, all results are normalized to a value of 1 at the wild-type length of 10 microns. Parameter values for the wild-type case are those previously derived from experimental measurements (Marshall and Rosenbaum, 2001). (**D**) Autocovariance of *lf1* (red) versus wild-type (blue) based on measurements of length fluctuations in living cells, showing higher mean squared fluctuations (based on autocovariance at zero lag) and slower decay for *lf* mutants, as predicted by the linear noise model. (E) IFT-coupled model for length fluctuations. Tubulin subunits (green) are brought to the end of the flagellum by anterograde IFT particles (blue) and deposited onto the tip, elongating the flagellum. Tubulin subunits disassembling from the tip are immediately brought back to the cell body by retrograde IFT particles (orange), shortening the flagellum. At steady state, elongation and shortening events occur at the same average rate but stochastic differences in the quantity of tubulin added or removed results in a net change in length of δ at a rate comparable to the frequency 1/τ of IFT trains arriving at or departing from the tip. (F) IFT-uncoupled model for length fluctuations. IFT maintains a pool of tubulin monomers near the tip, which undergo association or dissociation events at rates **K**^+^ and **K**^-^ respectively. Assembly rate **K**^+^ is length-dependent (see panel B), creating an effective restoring force. The resulting Ornstein-Uhlenbeck process defines the slope and asymptote of the mean-squared length change versus time plot, as indicated by the diagram, which can be directly compared to Figure 1B.

Clearly, average length can be increased either by increasing A, increasing P, or decreasing D. How would length-increasing parameter changes affect noise? To answer this question, we performed a small-signal linear noise analysis as derived in Materials and Methods, in order to ask how an intrinsic noise source within the flagellum would affect the uncorrelated variation in length between the two flagella. We do not know the ultimate source of intrinsic noise in this system, but we discuss possible sources in the Discussion section, below. Regardless of the source, growth rate fluctuations within the flagellum would cause the length to execute a one-dimensional random walk, as observed in **Figure 1B**, with the length-dependent assembly and disassembly rates created by the balance point model providing a restoring influence that returns the length back to its set point after any fluctuation, thus limiting the extent of fluctuations as shown by the plateau in **Figure 1B**. Modeling flagellar length fluctuations is thus analogous to the classical problem of modeling the Brownian motion of a particle attached to a spring. In our case the noise source is due to random variation in flagellar growth, for example due to stochastic fluctuation in IFT or in the addition/removal of tubulin, rather than thermal collisions, and the Hookean restoring force is a consequence of the fact that we use a linearized model to describe the rate of length change as a function of displacement from the steady-state solution. We further assume that the two flagella interact via competition for a shared limiting precursor pool P, as previously described (Ludington 2012). Under our assumptions, as derived in Materials and Methods, the variability of length differences between the two flagella in one cell is predicted to depend on model parameters as follows:

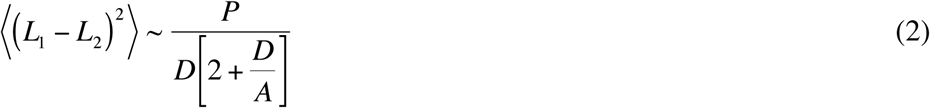

Results predicted by Equation 2 are plotted in **Figure 3C**. Note that any parameter change that increases flagellar length is predicted by Equation 2 to result in increased intrinsic noise, but changes in different parameters can have different relative effects on intrinsic noise. Changing either the cytoplasmic precursor pool size or the IFT efficacy have almost identical effects on noise, and in both cases the increase in the mean squared difference in the length of the two flagella (intrinsic noise) is linearly proportional to the increase in length, so a mutation that causes a doubling of average length should cause approximately a doubling of mean squared length differences. In contrast, changes to the disassembly rate D have a much larger, non-linear effect. It may thus be possible to predict which parameters of the model are affected by particular mutations, by determining the extent to which each mutation increases noise relative to average length. In any case, the first prediction of this model is that any parameter change that leads to an increase in length should also increase intrinsic noise in the balance-point length control system. This is consistent with our observations of increased length fluctuations in long flagella mutants and in cells treated with lithium (**Figure 2A**).

A second prediction of this model, derived in Materials and Methods, is that the effect of fluctuations will be less quickly damped out in mutants with long flagella. Indeed, the reason that the variance in length is predicted to increase in this model is that the effect of individual fluctuations should damp out more slowly in long-flagella mutants. This allows multiple fluctuations to add up to produce larger deviations from the average solution. Intuitively, we can see that fluctuations damp more slowly in long-flagella mutants due to the shape of the assembly versus length curve in the balance-point model (**Figure 3B**). Considering only small fluctuations, the rate at which any given fluctuation is corrected depends on the slope of the assembly-versus-length curve in the region of the steady-state solution, since the disassembly curve is a horizontal line. Since the assembly versus length curve is an hyperbola, its slope is a continuously decreasing function of length. For long flagella obtained via a decrease in disassembly rate, such mutants will have a decreased slope of their assembly curve at the steady state solution, thus fluctuations will persist for longer than wild-type. For long flagella obtained via an increase in either the precursor pool size or the efficacy of IFT, the length dependence of the assembly curve would be re-scaled to produce a lower slope at every value of length, again leading to slower damping of fluctuations. Thus, any parameter change that causes increased length would also cause fluctuations to be dampened more slowly.

In order to determine if the increased intrinsic noise in *lf1* mutants was due to larger or slower dynamic fluctuations, we analyzed fluctuations in individual *lf1* mutant cells using time-lapse microscopy. As shown in **Figure 3D**, we found that the fluctuation magnitude of individual flagella was approximately double in *lf1* compared to wild type (as judged by the autocovariance value for zero lag). This is the same order of magnitude increase as predicted by the theoretical noise model for mutations affecting either IFT or cytoplasmic precursor pool size, but would not be consistent with an alteration in disassembly rate. Moreover, the fluctuations were damped out more slowly in *lf1* compared to wild-type as judged by the decay of the autocovariance function, in full agreement with the model. A best fit exponential to the data of **Figure 3D** indicated a correlation time of 101 min for wild-type and 183 min for *lf1*. We conclude that the balance-point model can account for the nature of increased length fluctuation seen in *lf* mutants.

### Biological relevance of flagellar length variation

We and others have used flagellar length control as a paradigm for studying the general question of organelle size regulation. But is flagellar length physiologically relevant? Likewise, analysis of length fluctuation and variation can serve as a novel way to probe length control mechanisms. But does the cell care how much its flagella may vary in length? In order to investigate the biological significance of noise in flagellar length control, we asked whether uncorrelated noise (i.e., inequality in flagellar lengths) or correlated noise (i.e., cell to cell variation in the lengths of both flagella) affects flagella-driven motility. *Chlamydomonas* cells bend their two flagella in opposite directions in order to swim forward in a breast-stroke like motion. Inequality in lengths, as measured by uncorrelated noise, might thus be expected to prevent a cell from swimming forward effectively in a straight line. To test this prediction, we observed *Chlamydomonas* cells swimming using a high speed camera (**Video 1**), and measured the lengths of both flagella as well as swimming speed. To measure a range of different flagellar lengths, we analyzed motion of both wild-type and mutant cells with altered flagellar length but normal motile machinery, as discussed in Materials and Methods. As shown in **Figure 4A**, we found that optimal swimming occurred when both flagella were of approximately equal length and when their lengths were in the range 8-12 microns. Outside of this range, swimming speed decreased dramatically. The defects in swimming were different depending on the type of length variation. When both flagella were too long, the extra length of flagellum appeared to flail ineffectively through the media, presumably impeding forward progress (**Video 2**). This observation is consistent with an earlier report showing that *lf* mutants of Chlamydomonas display reduced swimming speed and beat frequency, with a reduction that correlates with the length of the flagella (Khona 2013). On the other hand, if both flagella were too short, they were unable to generate an asymmetrical bending motion. Consequently, the flagella just moved back and forth like flippers (**Video 3**) but could not create overall forward motion because symmetrical swimming movements do not produce motion at low Reynolds number (Purcell 1977). If the two flagella were of sufficiently unequal length, the cells swam in circles (**Video 4**). These results indicate that to the extent that a length control system may have evolved to ensure effective swimming, it must satisfy two goals: ensure equality of flagellar length within a cell, and ensure that the two flagella are in roughly the right average length range. Satisfying these two goals means that both correlated and uncorrelated variation would need to be constrained.

**Figure 4.**
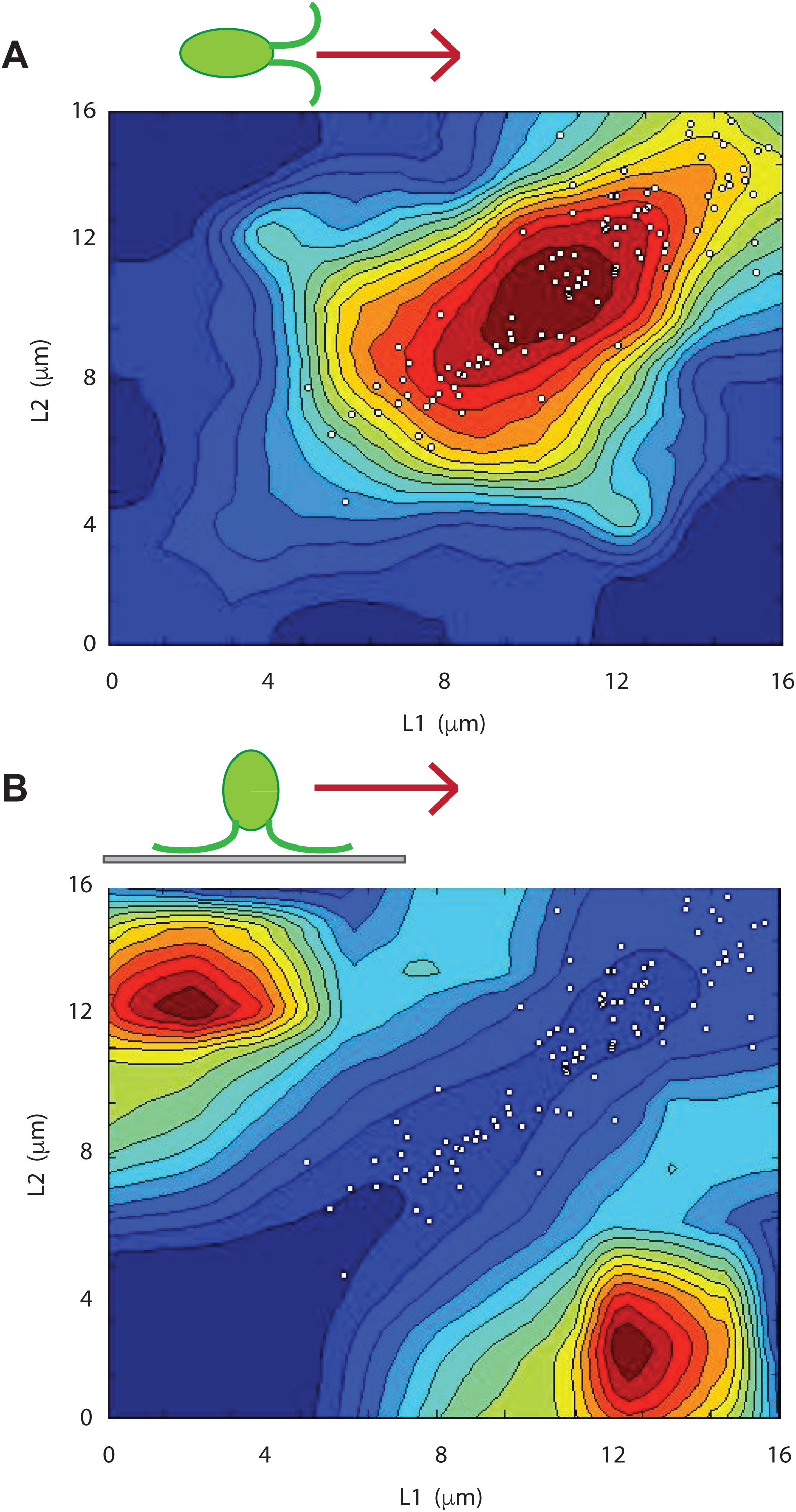
Noise affects fitness as judged by flagella-driven motility. (A) Contour plot of swimming speed (red fastest, blue slowest) versus the lengths of the two flagella. White dots signify lengths of flagella in wild-type cells superimposed on swimming speed distribution. Cells with the least uncorrelated variation (closest to diagonal) swim fastest in any given length range. (B) Contour plot of gliding speed (red fastest, blue slowest) versus the lengths of the two flagella. Cells with the most uncorrelated variation (farthest from diagonal) glide fastest in any given length range.

We next compared the actual distribution of flagellar lengths to the range of flagellar lengths that gives optimal swimming efficiency. As indicated by the dots in **Figure 4A**, wild-type cells have a joint distribution of lengths for their two flagella that corresponds to the optimal range of flagellar lengths for fast swimming. Thus, any increase in noise that would lead to variation outside this range would decrease swimming fitness. The joint length distribution in gametes matches the region of optimal swimming even more closely (**Supplemental Figure S2A**) due to the lower uncorrelated variation in gametes (see above).

Although flagella are used for swimming when cells are in suspension, they can also generate gliding motility when cells are adhered to a solid substrate (Bloodgood 1981). Gliding is an important type of motion for a soil-dwelling alga and one that is thought to possibly have evolved prior to bending-type flagellar motility required for swimming (Mitchell, 2007). Gliding involves motion of transmembrane proteins in the flagellar membrane adhering to the surface and then being pulled by a kinesin- and dynein-dependent motile process known as intraflagellar transport (IFT) (Shih et al., 2013; Collingridge 2013). Does flagellar length variation affect gliding motility in a similar way to its effects on swimming motility? To answer this question we imaged live cells during gliding using DIC microscopy and measured their flagellar lengths and gliding speeds, assessed as the distance the cell travelled over a fixed period of time (see Materials and Methods). The results of this analysis (**Figure 4B**) were strikingly different from those of the swimming speed analysis. Unlike swimming, which was fastest when the two flagella were of equal length, gliding was fastest when the two flagella were of unequal length (correlation coefficient between length difference and gliding velocity r=0.52, n=141, p<10^-7^). Gliding velocity correlated with the difference in the lengths of the two flagella, but not with the average of the two flagella (r=-0.014), confirming prior reports that overall flagellar length did not influence gliding speed (Bloodgood, 1981). Gliding thus appears to be sensitive to uncorrelated variation affecting the length of one flagellum versus the other, but not correlated variation affecting both flagella.

We noted that in gliding cells with unequal length flagella, cells always moved in the direction of the longer flagellum. We hypothesize that this behavior might reflect the tug-of-war nature of gliding motility (Shih et al., 2013; Collingridge 2013) such that one flagellum can only glide if the other flagellum is not gliding. In cells with equal length flagella, we would therefore expect to see the cell jitter back and forth rather than undergo long periods of directed movement, and indeed this is the case. As shown in **Supplemental Figure S2B**, we find that the frequency of reversal of gliding direction is highest in cells that have equal length flagella (r=-0.35 between length difference and reversal frequency, p=10^-5^). Cells in which one flagellum is much longer than the other show a low frequency of directional reversal, and tend to move at high speed in the direction of the longer flagellum. A more extreme form of this observation was previously reported by Bloodgood who observed that cells in which one flagellum had been sheared off always glided in the direction of the remaining flagellum and did not undergo any directional reversals (Bloodgood, 1981).

## Discussion

### Length control versus noise reduction

The results presented here show that the flagellar length control system in *Chlamydomonas reinhardtii* exhibits both intrinsic and extrinsic variation. The overall effect of these noise sources is that in a wild-type cell, the coefficient of variation (standard deviation in flagellar length divided by average flagellar length) is in the range 10-20%, which is comparable to the measured variability in length of vertebrate cilia (Wheatley and Bowser, 2002) and in the length of bacterial flagellar hooks (Koroyasu et al., 1998). While it is commonplace to compare cells and cellular structures with man-made machines, such a high level of size variability would not be tolerated in most manufacturing processes. It is therefore interesting to consider whether the level of noise that is observed indicates that cells cannot do any better, or alternatively that they do not need to. The close alignment between the distribution of flagellar lengths in wild-type cells and the length range for optimal swimming speed (**Figure 4A**) strongly suggests that the latter explanation is correct - there is little evolutionary pressure to reduce variability in flagellar length below a level that is tolerable for effective swimming. In fact, quite to the contrary, there may be an evolutionary pressure to maintain some degree of intrinsic noise to allow for better gliding motility. Variability in flagellar length may thus benefit a population of cells by giving the population as a whole more flexibility in dealing with changing environmental conditions. Gametes, which need to swim together in order to mate, showed reduced uncorrelated variation, possibly suggesting that they are further optimized for swimming at the expense of gliding.

### What is the noise source?

An important use of noise analysis is to discriminate among different feedback and control mechanisms for a system (Amir and Balaban, 2018). In some cases, analysis of fluctuations can be used to invalidate a model that otherwise accounts for the average behavior of a system. The present analysis of noise in flagellar length was aimed, in part, at attempting to invalidate our previous models for length control. The balance-point model has been shown to be able to account for the kinetics of flagellar regeneration (Marshall 2005), the dependence of flagellar length on flagellar number (Marshall 2005), and the ability of cells to equalize their flagellar lengths when one is severed (Marshall 2001; Ludington 2012; Hendel 2018). Here we find that our model can, at least qualitatively, account for the observation reported in **Figure 2** that long flagella mutants show increased uncorrelated variation (**Figure 3C**). Unfortunately, a quantitative comparison, for example between predicted and measured noise power spectra, would require more understanding of the actual noise source within the flagella. Currently, we do not know the source of the noise.

As with any chemical reaction, assembly and disassembly of tubulin subunits on the axonemal doublets is an inherently stochastic process and thus constitutes one unavoidable source of noise. Simulations of chemical master equations have been used to predict fluctuations in flagellar assembly and length at steady state based on the inherent stochasticity of tubulin monomer addition and removal (Banerjee 2020; Patra 2020). However, additional sources of variation may include stochastic fluctuations in cargo loading onto the IFT system (Craft 2015), transport of IFT trains through the flagellar pore (Dishinger 2010; Ludington 2014; Hendel 2018; Harris 2020), and fluctuation of IFT traffic due to traffic jams, motor pausing, and reversals (Bressloff 2006; Pinkoviezky 2014; Mijalkovic 2017; Tang 2019). For example, it has previously been proposed that entry of IFT particles into the flagellum may occur via an avalanche-like process, which naturally leads to a long-tailed distribution of IFT train sizes (Ludington 2013). Fluctuations in active disassembly would also contribute to noise.

Existing models of flagellar length dynamics fall into two classes depending on how they treat the role of intraflagellar transport. In tightly IFT-coupled models, it is assumed that tubulin is transported processively to the end of the flagellum by anterograde IFT particles, where it assembles onto the tip, elongating the flagellum. At the same time, tubulin subunits disassemble from the tip, possibly passively but more likely driven by microtubule depolymerizing enzymes. After disassembly, the tubulin binds to retrograde IFT particles and is trafficked back to the cell body. At steady state, assembly and disassembly rates balance. However, random variation in assembly or disassembly rates, for example due to avalanche-like behavior of IFT injection (Ludington 2013), will lead to stochastic differences in the assembly and/or disassembly steps. This type of tightly-coupled model has been used to simulate length fluctuations (Hendel 2018). We can model these differences as causing random increases or decreases of flagellar length in steps of size δ with an average time between steps of τ, the reciprocal of IFT arrival frequency.

In full-length flagella at steady state, the average frequency of IFT injection and arrival at the tip is approximately 1.25 train per second (Engel 2009). The slope of the mean-squared change in length versus time plot (Figure 1B) is 0.0014 μm^2^ / s. If we view the changes in flagellar length in terms of a random walk, with random steps of size δ taking place at a frequency of 1.25 steps per second (corresponding to a time between steps of τ = 0.8 seconds), we find, using the relation slope=δ^2^/τ (Berg 1983), that the size of the random steps per IFT train must be approximately 0.033 μm. Given that there are 233 protofilaments per cross section of flagellum, and each tubulin dimer added extends one protofilament by 8 nm, it would be necessary for each IFT train to carry approximately 960 tubulin dimers. At steady-state lengths, each IFT train consists of approximately 16 IFT particles (Pigino 2009; Vannuccini 2016). Thus, if random IFT train arrivals were driving the observed length fluctuations, it would be necessary for each IFT particle to carry on the order of 60 tubulins. This greatly exceeds the value of tubulins usually thought to associate with IFT particles (roughly 4 tubulins per particle; Bhogaraju 2014). We note that although the prior calculations of Bhogaraju et al. have found that flagellar growth rates were compatible with each IFT particle carrying just a few tubulins, it is important to recognize that processive growth requires far fewer assembly events in order to elongate the flagellum to a given length by slow and steady accretion onto the end. In contrast, the random walk dynamics of length in Figure 1B requires forward and backwards steps in length, that correspond to a larger number of tubulins per IFT particle compared to processive growth. There is no reason to think that the tubulin binding capacity per IFT particle increases once flagella reaches steady state. In fact, Craft and co-workers (Craft 2015) have reported that at steady-state length, many IFT trains do not appear to carry any tubulin at all. If a large fraction of IFT particles are not carrying any tubulin, then the remaining particles would need to carry even more tubulin subunits in order to account for the observed fluctuations. We conclude that statistical fluctuation in train injection through the flagellar port, or transit along the flagellum, or arrival at the tip, are unlikely to be able to explain the observed flagellar length fluctuations.

An alternative to the IFT-coupled model is an IFT-uncoupled model for length dynamics, in which IFT maintains a pool of tubulin monomers near the tip, but the addition or removal of tubulin at the tip takes place independently of the arrival or departure of IFT trains. In essence, the trains dump off their cargo, and then the cargo (tubulin) assembles according to its own kinetic processes. This type of model has been used to represent the dynamics of tubulin assembly in flagellar length control (Banerjee 2020; Patra 2020).

We represent tubulin monomers undergoing association or dissociation by rates **K**^+^ and **K**^-^ respectively. As indicated in Figure 1B, assembly is length-dependent, taking place at a rate k^+^(L) in units of microns per second, but disassembly is length-independent occurring at a rate k^-^ microns per second (this rate corresponds to the macroscopic disassembly rate D in our previous formulations of the balance-point model). These rates of length change are converted to rates of tubulin association and dissociation by assuming an effective monomer size δ, giving assembly and disassembly rates **K**^+^ = k^+^/δ and **K**^-^ = k^-^/ δ respectively. At steady state, **K**^+^= **K**^-^ = **K**. In this representation, the association and dissociation events lead to a random walk with a diffusion constant D_diff_= **K**_ss_δ, and the length-dependence of k leads to an effective restoring force given by the derivative of the rate of change in length with respect to the length, r=d(k^+^(L)-k^-^)/dL, evaluated at L=L_ss_. Given the model (Figure 3B) that assembly rate k^+^ is proportional to 1/L with a proportionality constant A, then r= A/L_ss_ ^2^ = k^+^/ L_ss_. Length fluctuations are thus governed by a random walk process constrained by a linear restoring term. For such a process, the plot of mean-squared length change versus time (Figure 1B) takes a well-defined form, diagrammed in Figure 3F, in which the mean squared length change increases linearly with time lag (for small time lag), with slope 2D_diff_, and then plateaus at twice the mean squared length (2 <L> = 2 D_diff_/r). As described above, the plot in Figure 1B shows a linear increase with a slope of 0.0014 μm^2^ / s. The plot reaches at plateau at approximately 2<L^2^>=14 μm^2^. Hence <L^2^> ∼ 7 μm^2^. We can relate observables <L^2^> and L to obtain the effective monomer size δ of the random process as follows:

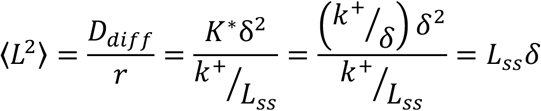

Using <L^2^> ∼ 7 μm^2^ from the plateau of fluctuation magnitudes in Figure 1B, and assuming that the average flagellar length is approximately L_ss_ =10 μm, we estimate the effective monomer size δ = 0.7 μm. Considering, as above, that each tubulin monomer is 8 nm long, and that there are 233 tubulins per cross section of the flagellum, the effective monomer size for the fluctuations of 0.7 μm would correspond to approximately 20,000 tubulin dimers. In order for this type of non-IFT coupled fluctuation model to work, the steps of the random walk cannot correspond to single tubulin dimer assembly or dissociation events. One possibility is that the random walk is driven by microtubule dynamic instability, in which periods of processive growth alternate with periods of processive shrinkage. This is analogous to the bursts of transcription that have been identified as an important source of noise in gene expression (Sanchez 2013).

### Using noise to test length control models

Another limitation of our currently modeling is that linear noise analysis of the type employed here is only strictly valid for very small fluctuations away from the steady state length. A more comprehensive approach would address the full nonlinear model. Several nonlinear stochastic models have recently been proposed for flagellar length dynamics (Fai 2019; Banerjee 2020; Patra 2020). Two of these studies analyzed models for flagellar length control using master equations to represent the inherently stochastic nature of biochemical reactions, which has the advantage of not requiring an extra “noise source” to be postulated. One study (Banerjee 2020) analyzed a model closely related to our balance point model, which assumes the 1/L dependence of IFT on length, while a second study (Patra 2020) analyzed a more elaborate model in which IFT injection is regulated by a time-of-flight mechanism. A third recent study describes a model for length regulation based on length-dependent transport of disassembly factors (Fai 2019), which is distinct from our current model which assumes length-independent disassembly. Analysis of this model, in which both IFT entry and the random walk of returning IFT motors are explicitly modeled as random processes, allowed for prediction of fluctuations in length, with the result that predicted noise is much smaller than observed fluctuations, assuming that the source of noise are addition and removal of single tubulins (Datta et al., 2020). Similarly, explicit modeling of IFT kinesin motor diffusive return showed that diffusion could produce a length-dependence that leads to a stable steady state length, and in combination with an avalanche-like entry of accumulated IFT particles at the flagellar base, could produce fluctuations in length.

In all of these cases, simulations indicate a fluctuation magnitude that could, with suitable choice of parameters, be made roughly comparable to what we have observed. As discussed in the previous section, many of these types of models require a very large “effective monomer” size, which may require a careful interpretation of this parameter in terms of actual molecular components or processes. An important future goal is to determine whether quantitative predictions from these models, such as the fluctuation power spectrum or the dependence of fluctuation magnitude on length, agree with the observations we have reported here.

Finally, we note that length control models have been proposed for other linear structures in the cell (Naoz 2008; Orly 2014; Mohapatra 2017), and we believe that noise analysis such as that reported here could be a productive way to probe these other models as well.

## Conclusions

Our results demonstrate that concepts of intrinsic and extrinsic noise are directly applicable to biologically relevant structure and function at the level of organelles, influence cell fitness, and are under genetic control.

## Materials and Methods

### Strains, media, and imaging

All strains were obtained from the Chlamydomonas Resource Center (University of Minnesota, St. Paul, MN), with the exception of *lf4* mutant strain V13 which was provided by Gregory Pazour, UMASS Medical Center. The *lf4* mutation was confirmed in this strain by PCR (data not shown). For asynchronous culture, cells were grown in 2mL cultures in TAP media (Harris, 1989) under continuous illumination. Cultures arrested in G1 were obtained by growing cells in M1 media for 2 days in continuous illumination and then switching them to continuous darkness for 24 hours. Gametes were grown overnight in M-N media.

To measure flagellar length in fixed cells, cells were fixed in 1% glutaraldehyde, and imaged using DIC optics with an Olympus 60x air lens and an air condenser on a DeltaVision 3D microscopy system. Three dimensional data was collected using a 0.2 μm step in the Z-axis. Lengths were then measured by tracing the flagella in three dimensions using the DeltaVision distance measuring function with the multi-segment length calculation method.

To measure flagellar length fluctuations in live cells, cultures were grown in TAP media at 21°C in constant light on a roller drum. For embedding live cells, 1% w/v agarose was melted in TAP media and cooled to 41°C. A square of Vaseline was made on a glass slide, then 5μL of culture was mixed with 20μL agarose in TAP within the square. The slide was then inverted over a coverslip, compressing the agarose in TAP into a flat, square pad. The cells were allowed to recover at room temperature in light for 2 hours to overnight, then imaged. The slide was imaged using DIC microscopy on a Deltavision microscopy system, at room temperature in ambient light. Embedded cells were imaged using a 100x oil immersion lens (NA 1.40 PlanApo) with a z step size of 0.2 microns. One z stack was taken through the entire cell every 10 minutes for 2 hours. The length of the flagella was measured using Deltavision software.

### Measuring flagellar length in live cells before and after pH shock

Two different apparatus were used separately o image flagella of individual *Chlamydomonas* cells during and after pH shock. In the first approach (**Figure 1E, S1A,B**), we used the CellASIC ONIX Microfluidic System (EMD Millipore, Hayward, CA). Cultured cells were loaded into the Microfluidic Chlamydomonas Trap Plate (C04A-01, EMD Millipore) and imaged on an inverted microscope (Ti-E Microscope, Nikon, Tokyo, Japan) with a 40× objective (Plan Fluor, 0.75 NA, Nikon) and an sCMOS camera (ORCA-Flash4.0, Hamamatsu Photonics, Hamamatsu, Japan) at 25°C. Cells were held in the microfluidic chamber in the plate by perfusing with TAP media at 5 psi and imaged by differential interference contrast (DIC) with 5 z-stacks at 0.9 µm intervals. To remove flagella from cells, we performed the pH shock method in the microfluidic chamber. Cells in the microfluidic chamber were deflagellated by perfusing with TAP media (pH 4.5, adjusted with acetic acid) for 1 minute, then neutralized by perfusing with TAP media (pH 9.0, adjusted with potassium hydroxide) for 30 seconds. After neutralization, the chamber was perfused with normal TAP media (pH 7.0) during the imaging. Cells were imaged every 10 minutes for 2 hours after pH shock deflagellation. Flagella were measured by hand-tracing in ImageJ (NIH, Bethesda, MA).

The long length of tubing in the CellASIC system raised concerns about the speed of solution exchange. We therefore implemented a second approach, by designing and fabricating a microfluidic device based on a serpentine channel (**Supplementary Figure S1C**) designed to allow rapid solution exchange while trapping cells. A single device is capable of immobilizing approximately 100 cells to a narrow z-plane (15 μm). The trapped cells are stable for many hours (>24), undergo cell division, and regenerate flagella. Unlike the CellASIC device, the serpentine channels are not suitable for imaging by DIC because flagella often lie near the channel wall and are thus difficult to image. We therefore used fluorescent imaging of cells expressing a flagellar marker. FAP20GFP (Yanagisawa et al,. 2014) cells were synchronized via a 14/10 light/dark cycle and hand injected using a syringe and a tube into the device until all traps were occupied. Constant flow was then established by attaching a syringe pump with normal and low-pH M1 media to the loaded device. A valve (Chrom Tech V-100D) was used to switch from normal to low-pH media with minimal lag time (**Supplemental Figure S1D**). The loaded device was mounted on a custom built OMX microscope (Dobbie et al,. 2011) with a 100X objective lens. An initial 3D-measurement (16 μm z-stack, axial spacing of 0.2 μm) was taken of flagellar length in the GFP channel (65μW @ 455nm at the sample). Cells were then pH shocked via switching to low-pH media for 60 seconds and returning to normal-pH media (**Supplementary Figure S1D**). Two hours after pH shock, cells were imaged again in the GFP channel for a final 3D length measurement. Length measurements were made by hand in ImageJ. XY and Z measurements were combined to compute overall length (**Supplementary Figure S1E,F**).

### Computing correlated and uncorrelated variation

We decompose the total variation in length into correlated and uncorrelated components following the relations for extrinsic and intrinsic noise described in dual-reporter gene expression noise studies (Elowitz et al., 2002). We do this in a purely operational sense by treating the two flagellar length measurements from each cell as though they were two fluorescence intensity measurements in a dual-reporter gene expression noise protocol, and computing the two components as follows:

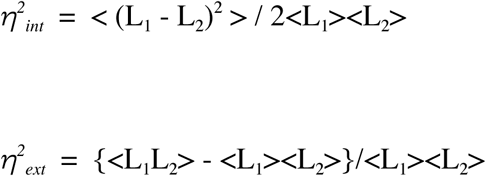

For each given cell measurement, we arbitrarily defined L_1_ as referring to the length of the flagellum whose base was closest to the left edge of the field of view.

### Estimation of measurement noise

We used time-lapse imaging to obtain two separate estimates for measurement error. First, as plotted by the black data-points in **Figure 1B**, we imaged cells fixed in glutaraldehyde at multiple sequential time points and then calculated the mean squared difference in length. As expected for measurement errors that are uncorrelated with each other, the slope of the best fit line to this data was less than 5×10^-5^, showing that the difference in measured length was independent of time lag. The average value of the mean squared difference in length was 0.0438 square microns, corresponding to an average measurement error of 0.2 microns.

As an alternative measure, we analyzed the fluctuations in live cells and calculated the mean squared change in difference between L_1_ and L_2_, and plotted this versus time. This plot gave a roughly linear behavior for the first several time points. We then fitted a straight line to the first ten time points and obtained a Y intercept of 0.597 microns squared. For a one-dimensional random walk (such as that executed by the difference in length between the two flagella), the Y intercept of this plot should correspond to 4 times the mean squared measurement error in each measurement. We thereby obtain an estimated measurement error of 0.39 microns. We thus obtain two independent estimates of measurement error, both of which are on the same order of magnitude as the XYZ voxel edge length, much smaller than the observed variations in length between flagella.

### Small-signal linear noise analysis of balance-point length control model

We have previously described a simple model for length control of cilia and flagella that has been termed the balance-point model. This model is based on observations that the axonemal microtubules undergo continuous turnover at their plus-ends, with disassembly taking place at a constant rate regardless of length, and assembly taking place at a rate limited by IFT (Marshall and Rosenbaum, 2001). A key component of the model is the hypothesis that intraflagellar transport is length-dependent. This dependency arises because the rate at which IFT particles enter the flagellum is proportional to 1/L (Marshall and Rosenbaum, 2001; Marshall et al., 2005; Engel 2009; Ludington 2013). The mechanistic reason for the 1/L dependence of IFT entry on length is not currently understood, although several models have been proposed (Ludington 2015; Ishikawa 2017b; Hendel 2018). Under the assumption that assembly is rate-limited by transport, it follows that the assembly rate is proportional to 1/L while measurements show that the disassembly rate is independent of length (Marshall and Rosenbaum 2001). Therefore, the assembly rate versus length curve will intersect the disassembly versus length curve at a unique value of the length, which reflects the steady state length of the flagella. The model presented here ignored a number of potential complications including the possibility that tubulin loading onto IFT particles may be regulated as a function of length (Craft 2015) and the fact that precursor production inside the cell body is actively regulated as a function of flagellar dynamics.

Following our previous formulation of the balance-point length control model, we define three parameters, A, P, and D. Parameter A describes the efficacy of intraflagellar transport and encapsulates the speed of the IFT particles, the number of IFT particles in a flagellum (a quantity known to be independent of length), and the cargo carrying-capacity of the particles. The parameter P describes the total pool of flagellar protein in a cell, including protein incorporated into the two flagella as well as unincorporated precursor stored in the cytoplasm (Rosenbaum et al., 1969). Parameter D describes the rate of flagellar disassembly. We have previously estimated values for these parameters in wild-type cells: D ∼ 0.22 μm/min (measured from the rate of shortening in *fla10* mutant cells), A ∼ 0.11 (this is a unit-less quantity), and P ∼ 40 μm (measured from the ratio of flagellar lengths before and after regeneration in the presence of cycloheximide (Marshall and Rosenbaum, 2001)). Note that the pool size is expressed in units of length, the conversion factor being the quantity of protein required to assemble a 1 μm segment of the axoneme.

The balance-point model can be encapsulated by a pair of coupled differential equations, one for each of the two flagellar lengths L_1_ and L_2_:

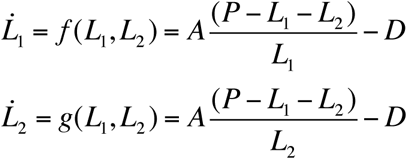

for which the steady state solution is:

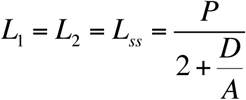

With the estimated parameter values given above, this predicts a steady state length of 10 μm. Using this model, we would like to predict how changes in the three parameters would affect noise suppression by the system, in order to ask whether parameter changes that increase the steady-state length would be predicted to increase noise. To do this, we employ a linear noise analysis by modeling the response to small perturbations in length.

We begin our noise analysis by introducing two variables:

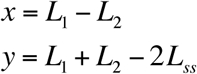

The new variables x and y correspond to the differences in flagellar length within one cell and the cell-to-cell variation in average length, respectively, that will result from a noise source applied to one of the two flagella, i.e., the response to an intrinsic noise source in one flagellum. We next consider the effect of applying a small perturbation to L_1_ at time t=0. Initially this perturbation would alter both x and y, but then over time the system will restore the flagellar lengths to their steady-state values. To determine how fast this restoration will occur, we linearize the system, as follows. First, we note that x and y can be expressed in terms of the deviations from the steady state in L_1_ and L_2_ as follows:

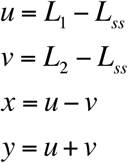

Assuming the perturbations are small, u and v will be close to zero. The rate of change of the deviations u and v is approximated by the Jacobian:

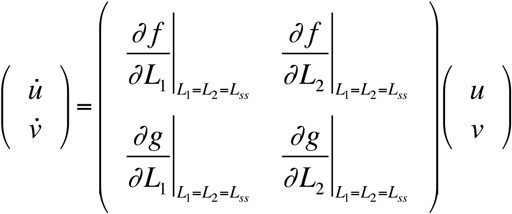

Evaluating the partial derivatives around the steady-state value L_1_=L_2_=L_ss_ we obtain the linearized system:

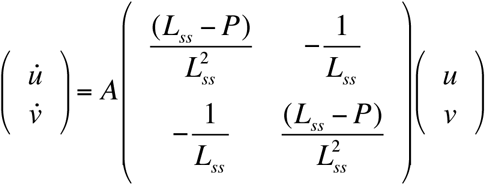

We then express the behavior of x and y in the linear approximation according to

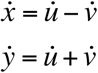

This substitution yields, finally, a pair of uncoupled differential equations for x and y:

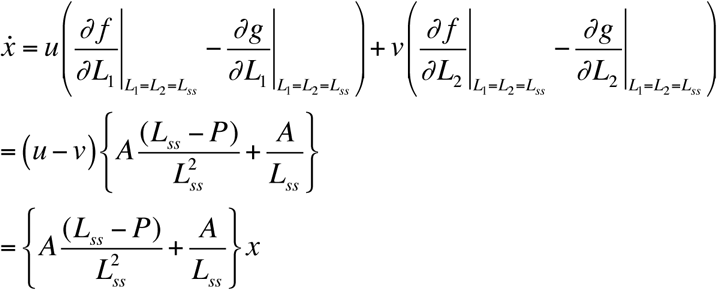

and

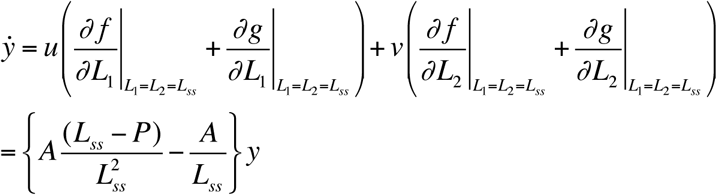

we define constants:

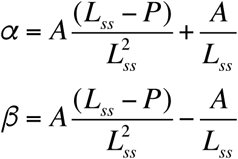

hence:

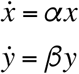

which, by inspection, have exponential functions as their solutions:

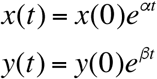

We now consider what happens if we add random fluctuations to the system. We model intrinsic fluctuations by adding a noise term to the equation governing x, thus:

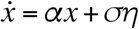

where σ denotes the magnitude of the fluctuations (in microns) and η denotes Gaussian white noise with zero mean and variance 1. The stationary solution to this equation (Honerkamp, 1994; Van Kampen, 1992) has a mean of 0 and a mean squared value of:

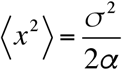

We assume that σ is the same for wild type and mutant cells. Hence the mean squared difference in length between the two flagella should be proportional to 1/α. Substituting the value of α derived above and then using the value of L_ss_ in terms of the three parameters A, D, and P, we obtain

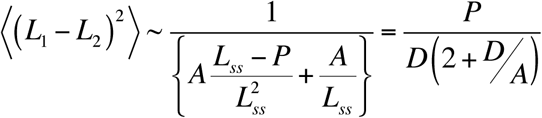

Which is given as Equation (2) in the Results section.

### Measurement of swimming speed versus flagellar length by high-speed video

To obtain simultaneous measurements of swimming speed and flagellar lengths for **Figure 4A** we mounted cells between a slide and coverslip supported by a 1 mm thick Vaseline ring, and imaged the cells on a Zeiss Axiovert 200M microscope with a 40X air objective lens using DIC optics and an infrared-blocking filter, collecting data with a Phantom MiroEx4-1024MM video camera at a frame rate of 1000 fps, exposure time 998 μSec. Cells were imaged at a dilution that ensured at most one or two cells per field of view. Following collection of each dataset, individual cells were manually tracked for 10 cycles of flagellar beating, marking the position of the cell at the beginning and end of this period. The difference in position divided by the elapsed time was taken as the average swimming speed. Since our data collection was only two-dimensional, many cells swam out of focus during the imaging and those images were discarded. Only cells for which the flagella remained in focus through the entire 10 beat cycles were used for distance measurements. In order to cover a wider range of lengths, we measured both wild-type cells and lf1 mutant cells, which frequently have unequal lengths and which span a range of short and long lengths. Swimming speed for lf1 mutants whose flagella are in the wild-type range are not statistically different from speeds of wild-type cells. Distance measurements of swimming and length measurement of flagella were performed using the Phantom Miro software.

### Measurement of gliding versus flagellar length

Cells were grown in TAP media and loaded onto a coverslip within a 2 mm thick Vaseline ring and inverted over a slide. Cells were imaged using a 20x objective with DIC optics. 25 images were collected at a rate of 1 image every 10 seconds. The total distance travelled by a cell was then calculated as the distance between the start and end point of the time series and the velocity calculated by dividing this distance by the total data collection time. The number of directional reversals for each cell during the entire time-course was determined by visual inspection.

## Acknowledgments

We thank Yee-Hung Mark Chan, Hiroaki Ishikawa, Tatyana Makushok, and Mark Slabodnick for careful reading of the manuscript, as well as Gabor Balaszi, Bob Bloodgood, and Ilya Nemenman, for helpful discussions and suggestions. Experiments to test flagellar length after regeneration were based on initial proof of concept by Zhiyuan Li and William Ludington. This work was supported by NIH grants R01 GM097017 and R35 GM130327, and by the Center for Cellular Construction (NSF DBI-1548297). JK was supported by the National Science Foundation grants DMR-1610737 and MRSEC-1420382, and by the Simons Foundation.

## Author Contributions

DB: Apparatus development, experiments and analysis (Figures 1F and S1C-F) Made figures, Wrote paper

HI: Experiments and analysis (Figure 1E, S1A,B)

KW: Experiments (Figure 1B, 3D)

JK: Mathematical modeling (Figure 3F) Data analysis (Figure 1B)

WFM: Experiments and analysis (Figure 1A, D, F; Figure 2, Figure 4, Figure S2) Data analysis (Figure 1B,C, Figure 3D) Mathematical modeling (Figure 3C,E) Made figures, Wrote paper

---

**Supplemental Figure S1.**
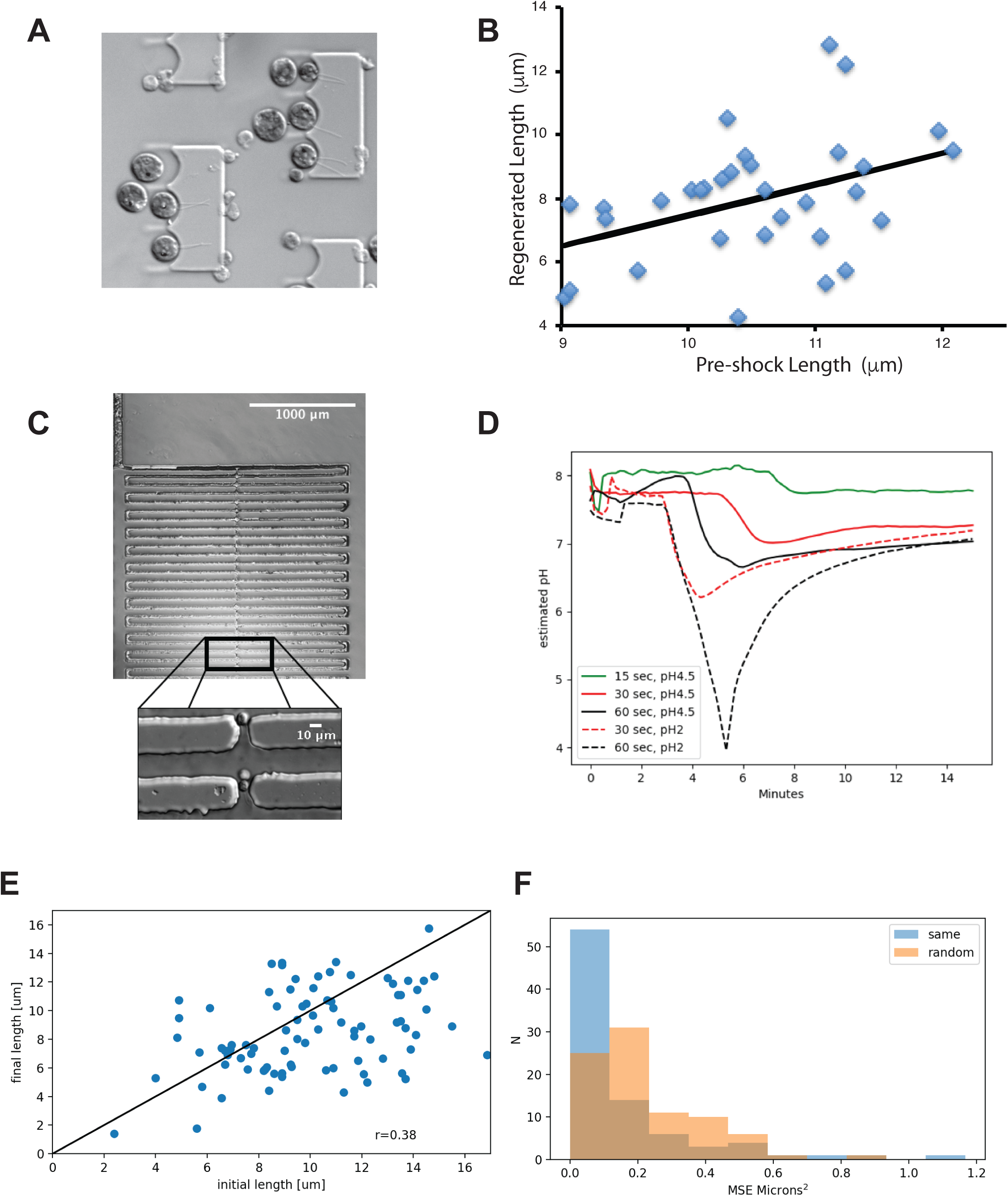
Additional data for Figure 1 about testing extrinsic noise by measuring flagellar length before and after regeneration. (**A**) Cells growing in commercial CellASIC chamber, based on a previously published design (Ludington 2012). (**B**) Length measurements before and after regeneration in *ptx1* mutant cells in CellASIC chamber (r=0.41; p=0.016; n=34). For this measurement, an outlier cell whose starting flagellar length was less than 7 microns was removed. (**C**) Design of new microfluidic trap for Chlamydomonas based on a serpentine channel. Lower image shows cells trapped by flow through holes in the serpentine channel. (**D**) pH change profile measured in new microfluidic device. (**E**) Length measurements before and after regeneration in wild-type cells in new fluidic device. To avoid potential confusion between the two flagella, both flagella lengths from one cell were averaged to compute correlation coefficients (r= 0.41; p=0.013; n=36). (**F**) As an additional test for cell-cell length variation, lengths before and after regeneration were compared between the same cell (blue) and randomly chosen cells (orange), and the mean squared difference in length recorded. The plot shows the histograms that result, indicating that comparison of flagellar lengths before vs. after regeneration in unrelated cells shows a greater difference than comparisons taken from the same cell.

**Supplemental Figure S2.**
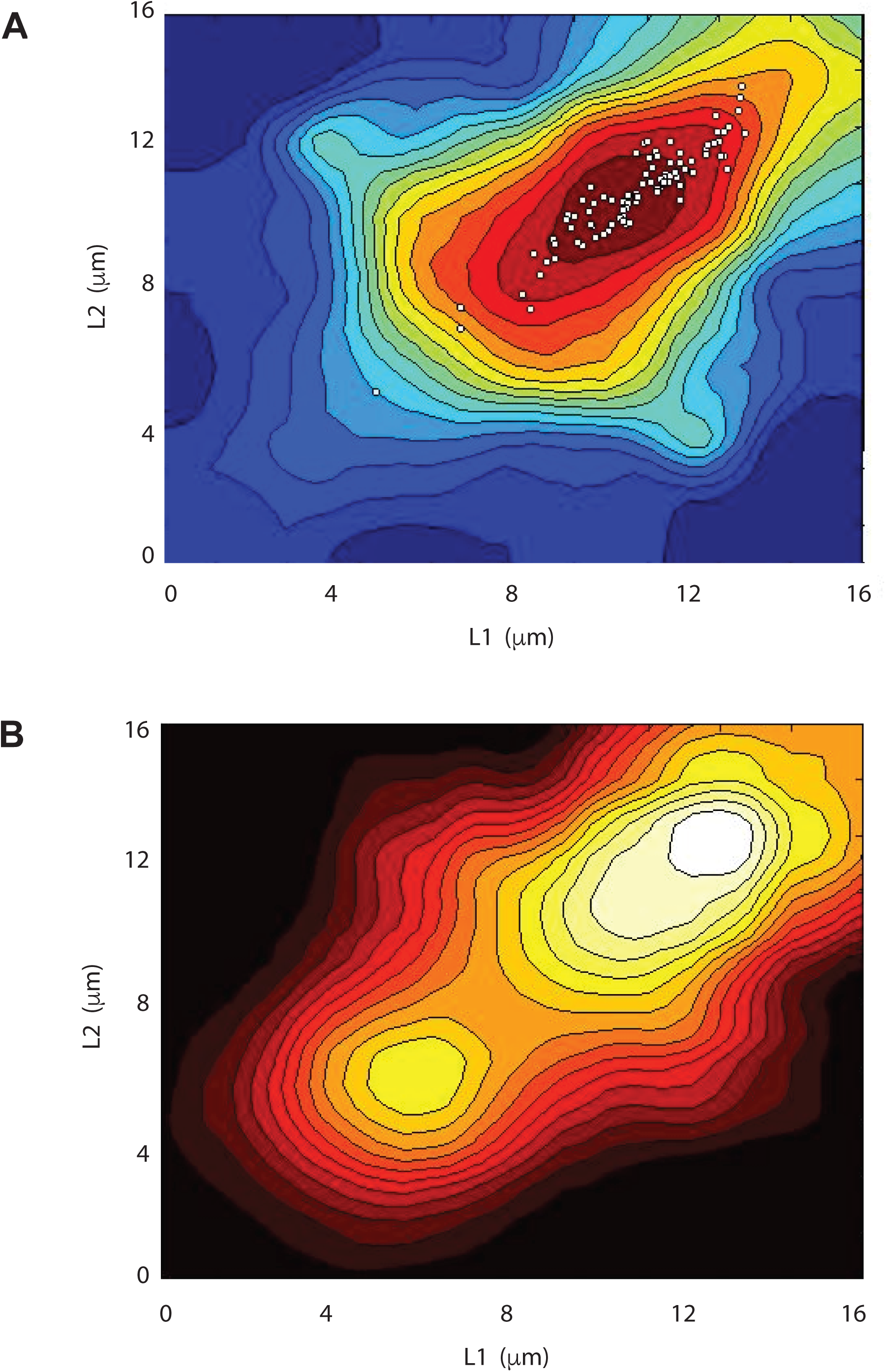
Additional data for Figure 4 regarding effects of flagellar length on motility. **(A)** Swimming speed versus flagellar length, with lengths of gametes superimposed as white boxes. Swimming speed data is the same as Figure 4A, but the gamete data-points illustrate that the length distribution of gamete flagella is better matched to the regime of optimal swimming. (**B**) Reversal frequency during gliding plotted versus flagellar lengths. White corresponds to maximum frequency of reversal, black minimum. The most reversals occur for cells with flagella of equal lengths, while increased length disparity correlates with decreased reversals.

**Video 1**. Wild-type *Chlamydomonas* cell imaged at 1000 frames per second, illustrating the breast-stroke motion of its two flagella.

**Video 2**. *Chlamydomonas lf1* cell with abnormally long flagella, showing uncoordinated motion when flagella are too long.

**Video 3.** *Chlamydomonas lf1* cell that has flagella half normal length, illustrating the inability to make forward progress with flagella that are too short to bend effectively.

**Video 4**. *Chlamydomonas lf1* cell with unequal length flagella (lower left quadrant, swimming in a circle).

